# La Crosse virus reassortants highlight genomic determinants of infection and pathogenesis

**DOI:** 10.1101/2024.03.11.584386

**Authors:** Nicole C. Rondeau, Sophie N. Spector, Sara A. Thannickal, Kenneth A. Stapleford

## Abstract

The genomic determinants that contribute to orthobunyavirus infection and pathogenesis are not well-defined. In this study, we harnessed the process of reassortment to understand which viral factors drive change in the replication and pathogenesis of La Crosse virus (LACV). We systematically reassorted the genomic segments of two genetically similar Lineage I LACV isolates into six unique reassortants. Despite the parental isolates having high levels of RNA and protein consensus, the reassortants demonstrate how minimal changes in RNA and protein structure can have significant changes in viral growth and reproduction *in vitro* in mammalian and insect models. We observed that swapping the S segment between isolates led to differences in replication and assembly resulting in one non-rescuable reassortant and one viable reassortant that exhibited an increase in viral growth dynamics. Switching the M segment led to changes in viral plaque phenotype and growth kinetics. L segment reassortants similarly differed in changes in viral growth dynamics. We further explored the M segment reassortants in a neonate mouse model and observed a role for the M segment in neuroinflammation and virulence. Through reassortment of the La Crosse virus genomic segments, we are able to further understand how genomic determinants of infection and pathogenesis operate in orthobunyaviruses. Future investigations will focus on identifying the specific molecular elements that govern the observed phenotypes *in vitro* and *in vivo*.

**Importance:** La Crosse virus is the leading cause of pediatric arboviral encephalitis in the United States, yet it is largely unknown how each of the three genomic segments contribute to pathogenesis and disease. Our study utilizes genomic reassortment between two similar Lineage I LACV isolates to understand genomic determinants for differences in infection and pathogenesis phenotypes in vitro and in vivo. By identifying roles for each segment in observed outcomes, we are able to plan further studies for molecular characterization of these phenotypes. Additionally, it is imperative to continue to characterize orthobunyavirus function since climate change will expand the range and prevalence of arthropod-borne diseases such as LACV in the United States.

## Introduction

The *Bunyavirales* order includes a diverse list of segmented RNA viruses capable of infecting varied hosts ranging from plants to vertebrates. The order is divided into multiple families with the majority of bunyaviruses being arthropod-borne viruses (arboviruses) spread to humans by mosquitoes, sandflies, and ticks. Importantly, many of these vector-borne bunyaviruses infect humans and livestock, causing devasting disease (1) (2) (3). However, there are no vaccines or antiviral therapies targeting bunyaviruses, highlighting the need to study these viruses in molecular detail.

La Crosse virus (LACV) is negative-sense segmented RNA orthobunyavirus transmitted to humans via infected mosquito bite (4) (5). LACV is endemic to the United States and is mainly found in the East North Central and Appalachian regions (6). A member of the California serogroup of orthobunyaviruses, it is related to other neuroinvasive human viruses found worldwide, including Jamestown Canyon virus, Inkoo virus and Tahyna virus (7). Although the majority of cases are asymptomatic and therefore underreported, LACV is the leading cause of pediatric arboviral encephalitis in the United States. Neuroinvasive disease can result in fatality or lifelong neurological sequalae, such as recurring seizures and cognitive deficits (8) (9) (10).

The LACV genome contains three single-stranded negative-sense RNA segments, which are referred to as the small (S), medium (M), and large (L) segments. The S segment encodes the nucleoprotein (N) and non-structural protein NSs, which has been shown to antagonize the mammalian type I interferon system (11) (12). The M segment encodes two glycoproteins (Gn and Gc), with Gc being a class II membrane fusion protein, as well as the non-structural protein NSm (13) (14) (15) (16) (17). Finally, The L segment encodes the RNA-dependent RNA polymerase (L) (18). While we understand which proteins are encoded by each segment, we do not completely understand how each protein and genomic segment, as well as discrete domains within these molecules, contribute to LACV replication and pathogenesis.

To identify and study LACV genomic determinants of replication and virulence, our study takes advantage of the two genetically similar Lineage I LACV strains that differ in plaque size and replication *in vitro* and *in vivo*. We created reassortants between these two isolates and found that S, M, and L segments each contributed to replication *in vitro* via unique phenotypes. Focusing on the M segment swap, we found that the M segment influenced plaque size, replication, and virulence and neuroinflammation *in vivo,* despite minor differences in viral titers in the brain. Interestingly, when we mapped the genetic differences between viral isolates, we found that the S segment only differed at the RNA level, whereas the M and L segments contain both protein and RNA changes. Taken together, these data identify critical roles for each LACV genome segment in replication and highlight important functions for the RNA and potentially protein sequence in LACV infection.

## Results

### Two genetically similar La Crosse virus strains confer differences in plaque size and viral growth kinetics *in vitro*

Previous studies have shown that different La Crosse virus (LACV) strains exhibited differences in viral growth kinetics and pathogenesis (19) (20). However, the molecular determinants driving these phenotypes are not well-defined. To begin characterizing these determinants, we acquired infectious clones of the Lineage I isolates, LACV_Human/1960_ (LACV_1960_) (11) and the LACV_Mosquito/1978_ (LACV_1978_) (21). We first looked to compare the genetic similarities between these two LACV isolates at the protein and RNA level (**Tables 1 and 2**). After aligning each segment, we found that the S segment proteins were 100% conserved, yet we found that there were slight amino acid differences between M and L segment protein pairs (**Table 1**). Interestingly, at the RNA level, we found that the S, M, and L segments all had variability between strains, with the M segment being the least conserved at 95.7% nucleotide identity (**Table 2**). This analysis led us to consider if these variations at the protein and/or RNA level would confer differences in the infectability and pathogenicity of LACV.

**Table 1:** Protein conservation between LACV strains.

|  | LACV<sub>Human/1960</sub><br>S segment | LACV<sub>Human/1960</sub><br>M segment | LACV<sub>Human/1960</sub><br>L segment |
| --- | --- | --- | --- |
| LACV<sub>Mosquito/1978</sub><br>S segment | 0 AA<br>100% |  |  |
| LACV<sub>Mosquito/1978</sub><br>M segment |  | 30 AA<br>97.9% |  |
| LACV<sub>Mosquito/1978</sub><br>L segment |  |  | 18 AA<br>99.2% |

*Aligned using ClustalW Multiple Sequence alignment*
*Number of amino acid differences indicated above consensus percentage*

**Table 2:** RNA conservation between LACV strains.

|  | LACV<sub>Human/1960</sub><br>S segment | LACV<sub>Human/1960</sub><br>M segment | LACV<sub>Human/1960</sub><br>L segment |
| --- | --- | --- | --- |
| LACV<sub>Mosquito/1978</sub><br>S segment | 22 bp<br>97.8% |  |  |
| LACV<sub>Mosquito/1978</sub><br>M segment |  | 195 bp<br>95.7% |  |
| LACV<sub>Mosquito/1978</sub><br>L segment |  |  | 284 bp<br>95.9% |

*Aligned using Smith-Waterman RNA alignment*
*Number of nucleotide differences indicated above consensus percentage*

Given a relatively high level of RNA and protein conservation, we wanted to first observe if there are any growth kinetic differences in insect or mammalian cells between our two wild-type strains. To generate virus, we transfected LACV_1960_ and LACV_1978_ S, M, and L segment plasmids into BHK BSR-T7 cells, where a T7 promoter drove expression of each plasmid. After passaging our viruses on Vero cells, we collected the viral supernatants and confirmed their sequences via Sanger sequencing. We then plaqued both strains on Vero cells to see whether they were able to produce infectious particles. Interestingly, we found a distinct plaque phenotype between both stocks, with the LACV_1978_ producing a much larger plaque size compared to the LACV_1960_ (**Figure 1A**).

**Figure 1:**
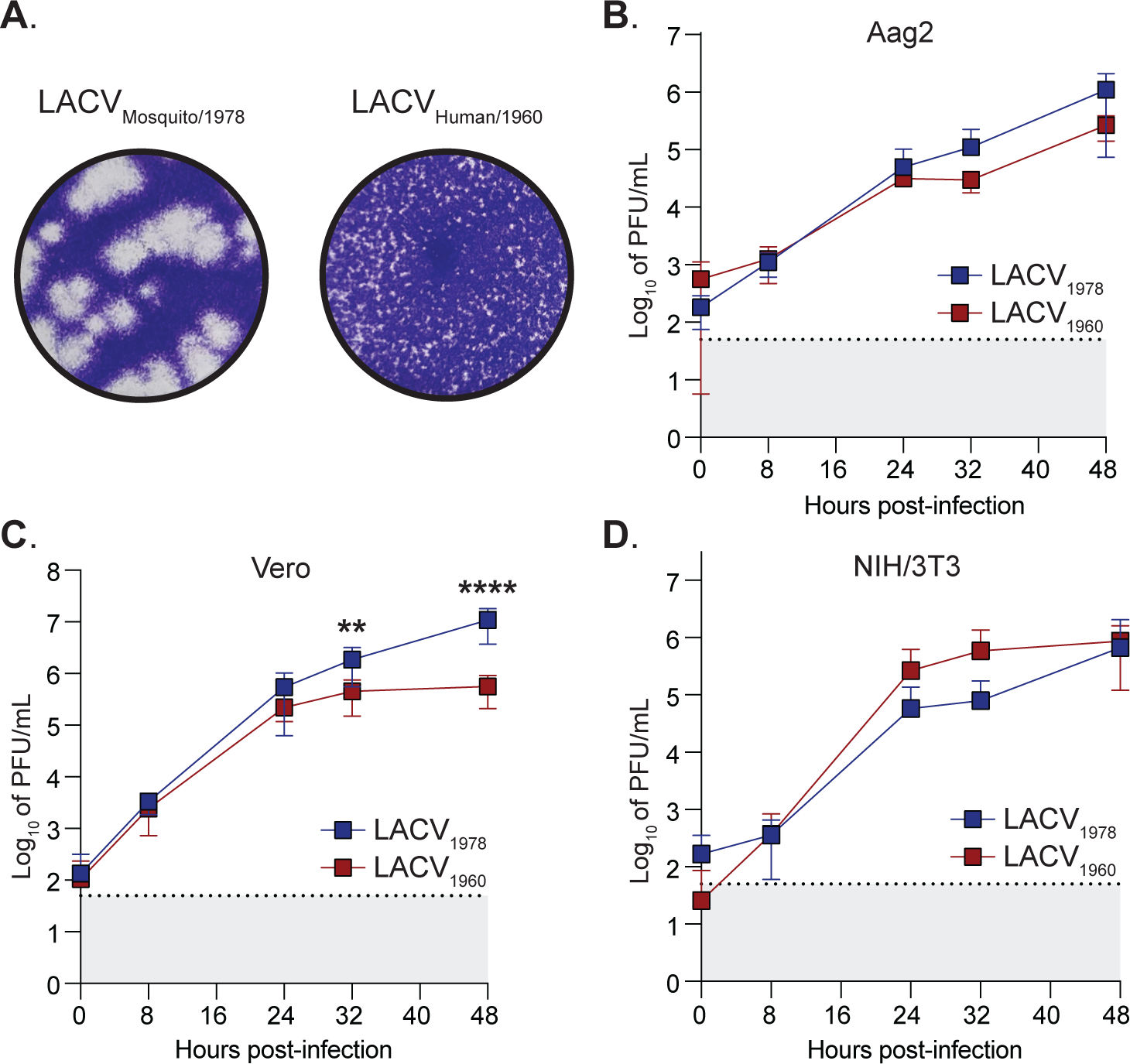
La Crosse virus WT strains exhibit differences in growth kinetics in mammalian and insect cells. (**A**) Representative plaque assays on Vero cells of both LACV_1978_ and LACV_1960_ WT strains. Plaques were fixed and developed after 4 days post-infection. (**B**) Aag2 cells, (**C**) Vero cells and (**D**) NIH/3T3 cells were infected at a MOI of 0.03 for insect cells and 0.1 for mammalian cells. After 1 hour, cells were washed with PBS and supernatants were collected at 0, 8, 24, 32, and 48 hours post-infection to quantify viral titers via plaque assay. The dotted line and gray shaded area represent the limit of detection (LOD). Data represent at least three independent trials with internal duplicates. Statistical significance was found via Mann-Whitney tests, with p-values representing **p < 0.01 and ****p < 0.0001. The mean and positive-value standard deviation (SD) are shown for all data.

To test for differences in growth kinetics, we infected Aag2 mosquito cells at a MOI of 0.03 (**Figure 1B**), and mammalian Vero (**Figure 1C**) and NIH/3T3 (**Figure 1D**) cells at a MOI 0.1. Supernatants were collected at timepoints 0, 8, 24, 32, and 48 hours post-infection and viral titers were quantified via plaque assay. We found that the LACV_1978_ strain has a significant growth advantage in Vero cells, and a slight advantage in our Aag2 insect cells after 32 hours post-infection. On the contrary, in NIH/3T3 cells, the LACV_1978_ strain showed attenuation in growth between 24 and 32 hours, but then matches the LACV_1960_ strain by 48 hours post-infection. These differences in growth kinetics allow us to hypothesize a potential host-cell specificity between our two strains as well as strain-specific genomic determinants that are influencing phenotypic differences.

### La Crosse virus S Segment reassortment results in differences in virus rescue and growth kinetics *in vitro*

Given the genetic similarities between these two strains, we hypothesized that we could use reassortment to begin identifying the genomic determinants driving the phenotypes in Figure 1. We systemically swapped one of the three genomic segments between our LACV isolates, LACV_1960_ and LACV_1978_, resulting in six unique reassortants with two genomic segments from one LACV WT isolate and one genomic segment from the other LACV WT isolate.

We began by making swaps between the S segments, creating the reassorted viruses: **S_1960_**M_1978_L_1978_ and **S_1978_**M_1960_L_1960_ (**Figure 2A**). Reassortment was accomplished via transfection, as described above with WT LACV_1960_ and LACV_1978_ viruses. Interestingly, the reassorted virus **S_1960_**M_1978_L_1978_ was not able to be rescued despite multiple rescue attempts (**Figure 2A**). However, the inverse reassortant, **S_1978_**M_1960_L_1960_, was viable and able to be propagated. These data suggest that the **S_1960_**M_1978_L_1978_ virus may contain a defect in viral assembly or packaging, despite 100% protein conservation.

**Figure 2:**
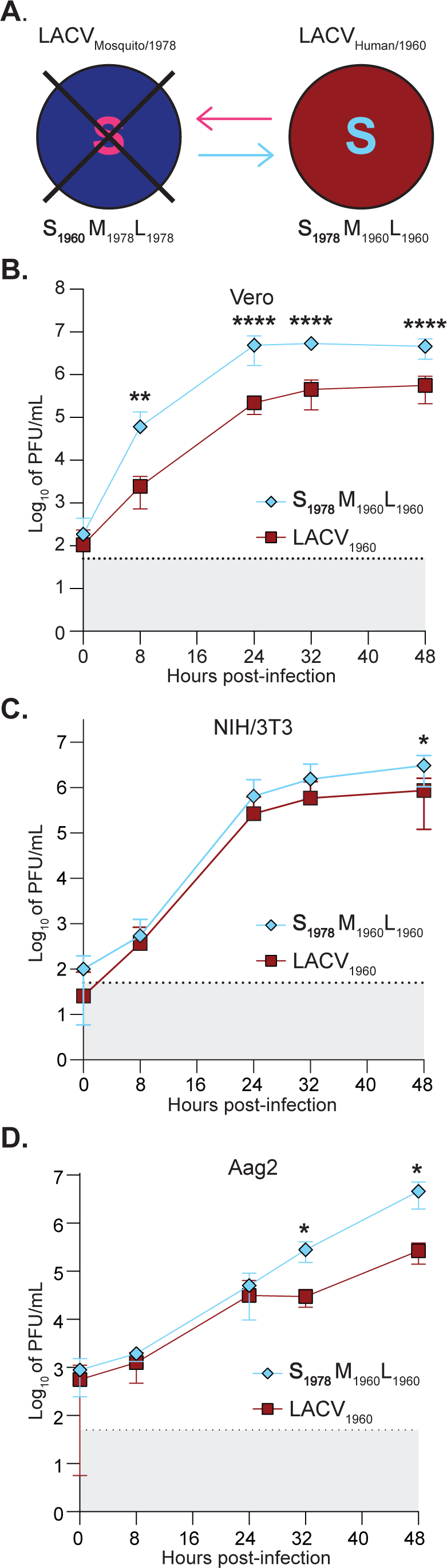
Reassortment of La Crosse virus S segment results in two distinct viability phenotypes. (**A**) Schematic indicating swap of S segment of LACV_1978_ and LACV_1960_ WT viruses to create reassortants: **S_1960_**M_1978_L_1978_ and **S_1978_**M_1960_L_1960_. Large X indicates failure to rescue virus. (**B**) Vero cells and **(C)** NIH/3T3 cells were infected with a MOI of 0.1 and (**D**) Aag2 cells were infected with a MOI of 0.03 of each virus. Following one hour, virus was removed and cells washed. Cells were incubated for 48 hours in complete media. At 0, 8, 24, 32, and 48 hours post-infection, supernatant was collected for plaque assay to determine infectious particle titers. Data represent at least two independent trials with internal duplicates. The dotted line and gray shaded area represent the limit of detection (LOD). A Mann-Whitney test was performed and statistical significance is shown (*p < 0.05, **p < 0.01, and ****p < 0.0001). The mean and positive-value standard deviation (SD) are shown for all data.

To determine if there are growth dynamic differences of **S_1978_**M_1960_L_1960_ in comparison to LACV_1960_, a multi-day growth curve was performed. Vero and NIH/3T3 cells were infected at a MOI of 0.1 while Aag2 cells were infected at a MOI of 0.03. Supernatant was then collected at multiple time points and infectious particle numbers were determined via plaque assay. We observed that the **S_1978_**M_1960_L_1960_ virus had a significant growth advantage over the WT LACV_1960_ virus in Vero cells early during infection (**Figure 2B**). In NIH/3T3 and Aag2 cells (**Figure 2C and D**), an eventual significant increase in viral titers of the **S_1978_**M_1960_L_1960_ virus when compared to WT LACV_1960_ at time points 32 and 48 hours post-infection was seen. Despite having near identical RNA sequences and identical amino acid sequences (**Table 1** and **2**), swapping the S segment between the two WT LACV isolates resulted in **S_1960_**M_1978_L_1978_ being non-rescuable while **S_1978_**M_1960_L_1960_ conferred a growth advantage in all cell lines.

### Swapping La Crosse virus M Segment changes plaque phenotype and viral titers *in vitro*

As previously mentioned, the two WT isolates used in this study, LACV_1960_ and LACV_1978_, have distinct viral plaque size phenotypes; LACV_1960_ results in small plaques while LACV_1978_ has a large plaque phenotype (**Figure 1A**). Given that the M segment of LACV is responsible for production of the glycoproteins Gc and Gn, which determine viral cell fusion and entry (22), we hypothesized that swapping the M segment between the two isolates would alter the plaque size phenotype. We reassorted the M segments between our two LACV WT isolates using the same method as before. Both reassortants, S_1978_**M_1960_**L_1978_ and S_1960_**M_1978_**L_1960_, generated rescuable viruses (**Figure 3A**). Swapping the M segment between LACV_1960_ and LACV_1978_ resulted in a change of the plaque size phenotype (**Figure 3B**). S_1978_**M_1960_**L_1978_ resembled LACV_1960_ with small plaques instead of having large plaques like its WT backbone, LACV_1978_. Inversely, S_1960_**M_1978_**L_1960_ exhibited large plaques like LACV_1978_ despite having two segments, S_1960_ and L_1960_, from LACV_1960_.

**Figure 3:**
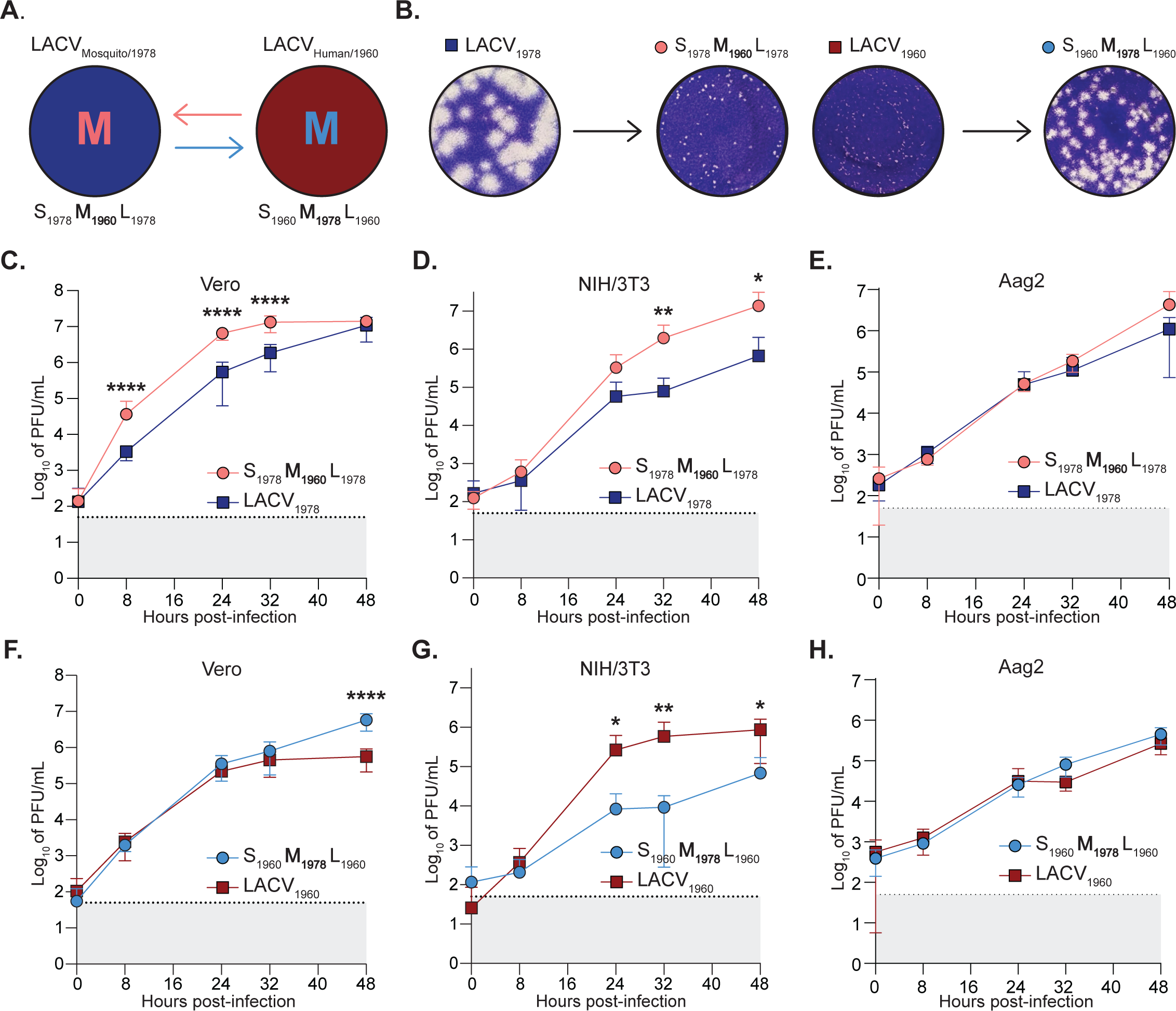
La Crosse virus M segment reassortment elicits plaque phenotype change and differences in viral replication. (**A**) Schematic showing swap of LACV_1978_ and LACV_1960_ WT virus M segments to create the reassorted viruses S_1978_**M_1960_**L_1978_ and S_1960_**M_1978_**L_1960_. (**B**) Representative plaque assays of LACV_1978_, S_1978_**M_1960_**L_1978_, LACV_1960_, and S_1960_**M_1978_**L_1960_, respectively. Plaques were fixed and developed 4 days post-infection. (**C** and **F**) Vero cells and (**D** and **G**) NIH/3T3 cells were infected with a MOI of 0.1 and (**E** and **H**) Aag2 cells were infected with a MOI of 0.03 of each virus. After one hour, virus was removed, cells were washed with PBS and then incubated in complete media. Supernatant was collected at 0, 8, 24, 32, 48 hours post-infection for plaque assay to determine infectious particle titers. Data represent at least two independent trials with internal duplicates. The dotted line and gray shaded area represent the limit of detection (LOD). A Mann-Whitney test was performed and statistical significance is shown (*p < 0.05, **p < 0.01, and ****p < 0.0001). The mean and positive-value standard deviation (SD) are shown for all data.

To address the role of the M segment in virus replication, we again performed growth curves in mammalian and mosquito cells. In Vero cells, S_1978_**M_1960_**L_1978_ had an initial significant growth advantage over LACV_1978_ before plateauing in growth at 32 hours post-infection, similar to what was seen with the S swap **S_1978_**M_1960_L_1960_ (**Figure 2B**). By hour 48, LACV_1978_ had produced an equivalent amount of virus as S_1978_M_1960_**L_1978_**(**Figure 3C**). Conversely, S_1960_**M_1978_**L_1960_ and LACV_1960_ grew at equivalent rates until 32 hours post-infection where the reassortant produced significantly greater amount of virus at 48 hours post-infection (**Figure 3F**). In NIH/3T3 cells, S_1978_**M_1960_**L_1978_ produced significantly more virus than LACV_1978_ by 32 hours post-infection when grown (**Figure 3D**). Interestingly, S_1960_**M_1978_**L_1960_ exhibited a significant reduction in virus production when compared to LACV_1960_ 24 hours post-infection when NIH/3T3 cells are infected (**Figure 3G**). Infection of Aag2 cells elicited no difference in virus production between reassortant viruses and WT viruses (**Figure 3E and H**). Taken together, these data indicate that the LACV M segment determines plaque phenotypes likely due to glycoprotein sequence differences between isolates. Additionally, these minute changes in sequence lead to changes in growth dynamics when compared to the wild-type isolates.

### L Segment reassortants mimic M segment reassortant growth phenotypes *in vitro*

The LACV L segment encodes the RNA-dependent RNA-polymerase and therefore plays an integral role in LACV replication (23). To understand how the L segment may influence virus replication, L segment reassortants, S_1960_M_1960_**L_1978_** and S_1978_M_1978_**L_1960_**, were made using the aforementioned protocol and growth curves were performed. In Vero cells, virus production did not differ between S_1978_M_1978_**L_1960_** and WT LACV_1978_ (**Figure 4B**). However, the opposite switch, S_1960_M_1960_**L_1978_**, had a significant growth advantage over WT LACV_1960_ at all time points (**Figure 4E**). This is similar to the reassortant S_1978_**M_1960_**L_1978_ which has the same combination of M and L segments (**Figure 3C**). No significant differences in viral growth dynamics between coupled reassorted and WT viruses was seen in infection of NIH/3T3 (**Figures 4C and F**). However, it should be noted that S_1978_M_1978_**L_1960_** virus produced less infectious virus than WT LACV_1978_ after 24 hours post-infection. The only other instance of a reassortant producing less virus than its paired WT virus is between S_1960_**M_1978_**L_1960_ and LACV_1960_ (**Figure 3G**). Both of these reassorted viruses contain the M segment from LACV_1978_ and the L segment from LACV_1960_ suggesting that this pairing may negatively affect infectious particle production. Finally, similar to what was seen for the M segment swap, there was no difference in viral growth dynamics between S_1978_M_1978_**L_1960_**and LACV_1978_ and S_1960_M_1960_**L_1978_**and LACV_1960_ when grown in Aag2 cells (**Figures 4D and G**). Taken together, L segment reassortant growth phenotypes mirror those of their counterpart M segment swaps, demonstrating a potential interaction between M and L segments that is not influenced by presence of the S segment.

**Figure 4:**
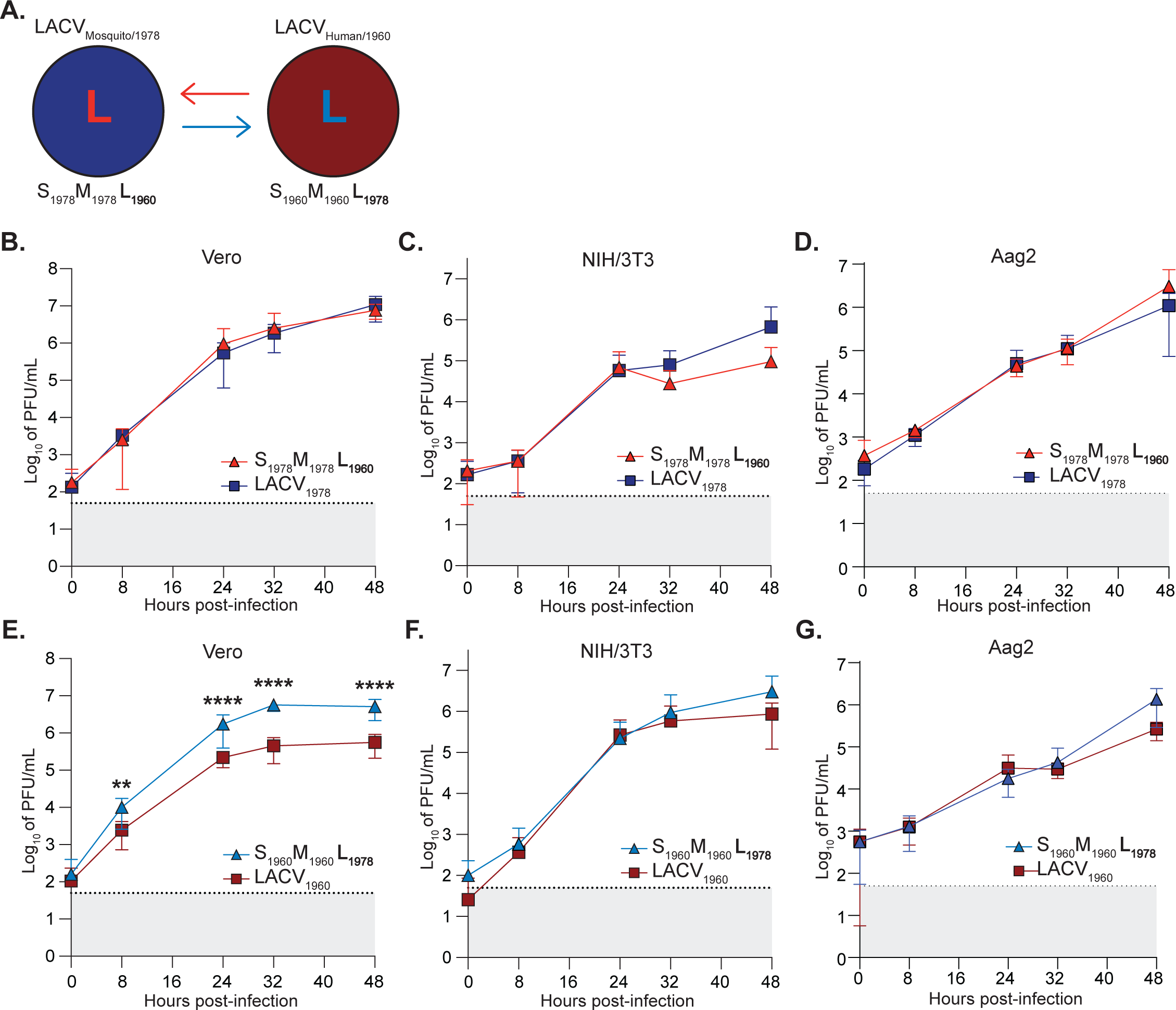
L segment reassortments mimic M segment reassortant growth phenotypes. (**A**) Schematic indicating reassortment of L segment of LACV_1978_ and LACV_1960_ WT viruses in order to create reassortants S_1978_M_1978_**L_1960_** and S_1960_M_1960_**L_1978_**. (**B** and **E**) Vero cells and (**C** and **F**) NIH/3T3 cells were infected with a MOI of 0.1 and (**D** and **G**) Aag2 cells were infected with a MOI of 0.03 of each virus for 1 hour. Cells were then washed in PBS and incubated in complete media for 48 hours. At 0, 8, 24, 32, and 48 hours post-infection, supernatant was collected for plaque assay to determine infectious particle titers. Data represent at least two independent trials with internal duplicates. The dotted line and gray shaded area represent the limit of detection (LOD). A Mann-Whitney test was performed and statistical significance is shown (**p < 0.01and ****p < 0.0001). The mean and positive-value standard deviation (SD) are shown for all data.

### M segment reassortants alter clinical outcomes and virulence *in vivo*

Previous studies have shown that the M segment of LACV and other orthobunyaviruses contributes to neuroinvasion and neuropathogenesis *in vivo* (24) (25) (26). However, we do not completely understand how the M segment contributes to disease. Therefore, we hypothesized we could use the M swap viruses to better understand how LACV replicates *in vivo*. Since La Crosse virus is the leading etiological agent of pediatric arboviral encephalitis in the United States and causes hospitalized instances of disease in humans less than 16 years old, the murine model we chose to study our LACV reassortants disease utilizes C57BL6/J neonatal mice (20). To test the neuroinvasive capacity of the M segments from our two LACV strains, we infected 6-day old mice with 50 PFU of S_1978_**M_1960_**L_1978_ or WT LACV_1978_ (**Figure 5A-C**) and 5-day old mice with 50 PFU of S_1960_**M_1978_**L_1960_ or WT LACV_1960_ (**Figure 5D-F**). At 3 days post-infection, neonates were observed for signs of disease and then assigned a clinical score (**Table 3**) based on the severity of their symptoms (**Figure 5A** and **D**). The pups were then euthanized and the brain was harvested for quantification of viral genomes via RT-qPCR (**Figure 5B** and **E**) and infectious titers via plaque assay (**Figure 5C** and **F**). While there are slight differences in viral titers between S_1978_**M_1960_**L_1978_ and LACV_1978_, we see a more drastic difference in the clinical scores of these pups, as the WT LACV_1978_ caused severe moribund disease symptoms that the M segment reassortant virus did not. In fact, infecting 5-day old mice with WT LACV_1978_ led to death within 3 days of infection (data not shown), which led us to include 6-day old mice in our study. By comparison, WT LACV_1960_ leads to moderate, but not severe, disease in 5-day old mice (**Figure 5D**). In line with this difference, the S_1960_**M_1978_**L_1960_ reassortant both had significantly higher titers and a difference in disease severity compared to LACV_1960_. Interestingly, the LACV_1960_ strain only had 40% of infected pups produce infectious titers, as compared to the other LACV_1978_ strain and two M segment reassortants, which had 100% of infected pups produce infectious titers. These results indicate that the M segment of the LACV_1978_ strain not only contributes to an increase in neurovirulence compared to the LACV_1960_, but also that the other LACV_1978_ segments contribute to severe disease as compared to the other strain and reassortants.

**Figure 5:**
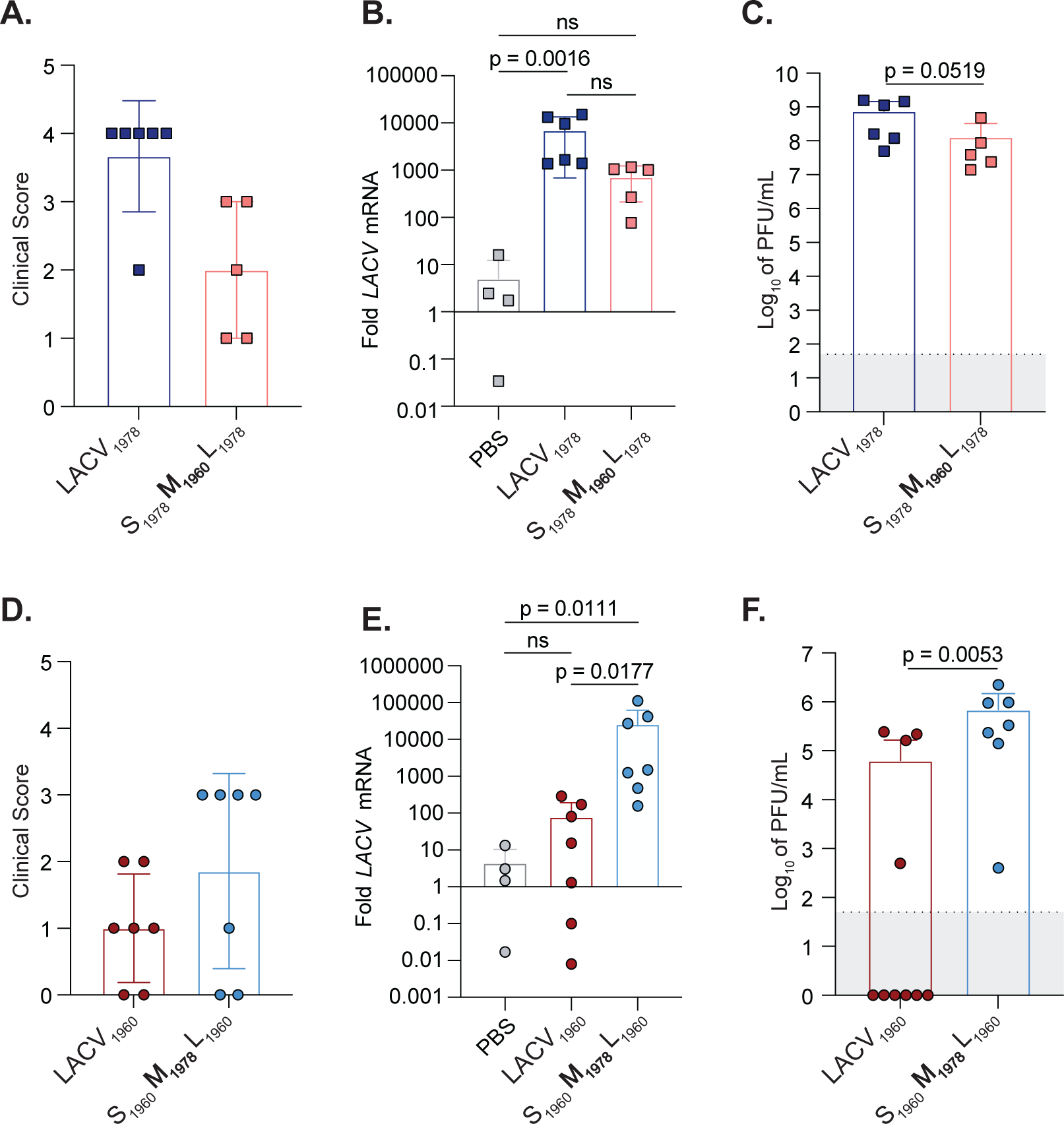
LACV_1978_ M segment contributes to neurovirulence in neonatal mice. (**A-C**) 6-day old (**D-F**) or 5-day old C57BL/6J mice were infected with 50 PFU of either a LACV M reassortant or its WT parent strain. After 3 days post-infection, pups were examined for signs of disease and assigned a clinical score (**Table 3**) (**A** and **D**) before euthanizing and harvesting the brain to determine viral genomes via qPCR (**B** and **E**) and viral titers via plaque assay (**C** and **F**). Data represent at least two independent trials, with n ≥ 4 mice. PBS controls shown are pooled from 5-and 6-day old mice. A Mann-Whitney test was performed and p-values are as shown (ns = non-significant). The mean and positive-value standard deviation (SD) are shown for all data.

**Table 3:**
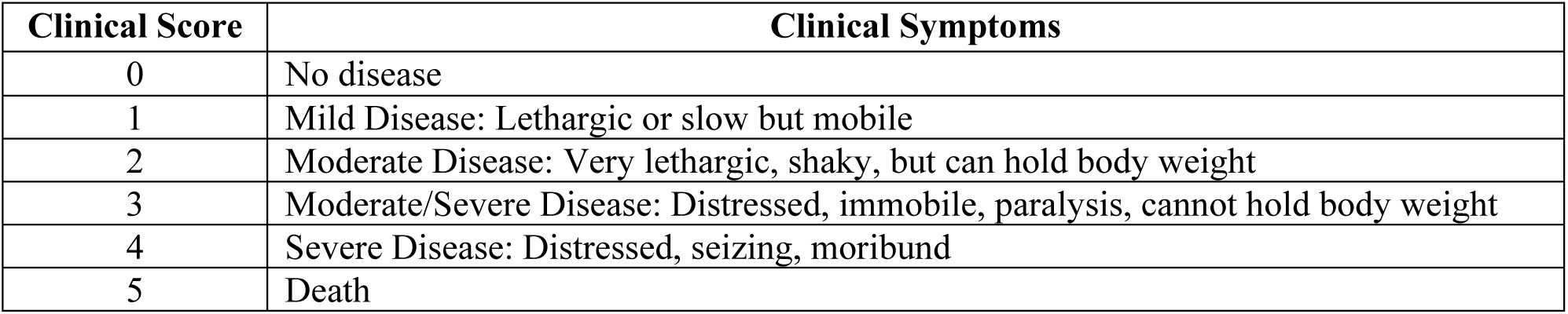
Clinical Score Guidelines.

| Clinical Score | Clinical Symptoms |
| --- | --- |
| 0 | No disease |
| 1 | Mild Disease: Lethargic or slow but mobile |
| 2 | Moderate Disease: Very lethargic, shaky, but can hold body weight |
| 3 | Moderate/Severe Disease: Distressed, immobile, paralysis, cannot hold body weight |
| 4 | Severe Disease: Distressed, seizing, moribund |
| 5 | Death |

### La Crosse virus M segment contributes to neuroinflammation *in vivo*

After observing notable differences in clinical outcomes, we next wanted to evaluate inflammatory markers in pup brains infected with LACV M segment reassortants. Because infectious titers were similar between LACV_1978_ and S_1978_**M_1960_**L_1978_ despite notable differences in clinical outcomes (**Figure 5C**), we suspected that alterations in neuropathogenesis and levels of inflammation may be mediating differences in disease outcome. Therefore, we quantified the relative expression of a panel of innate immune markers in the brain of infected mice by RT-qPCR (**Figure 6 and 7**). All gene-expression values are shown as fold change compared to PBS-infected controls, for which we pooled 5-and 6-day old control mice. First, we evaluated both type I interferon (IFN) (*Ifn-α, Ifn-β*) and type II IFN (*Ifn-γ*) RNA expression. Type I IFN is a well characterized mediator of LACV neuroinvasion and neuropathogenesis, and has been shown to be responsible for protecting against less neuroinvasive California serogroup viruses in addition to protecting adult mice from neurological LACV disease (27) (28). *Ifn-γ* is another antiviral cytokine that mediates protection from several RNA and DNA viruses in the CNS, and has been shown to be upregulated in neuron/astrocyte co-cultures infected with LACV as early as 24 hours (29). Comparing WT LACV_1978_ and the S_1978_**M_1960_**L_1978_ M segment reassortant, we found that the M_1960_ segment reduced levels of *Ifn-α* and *Ifn-β* RNA expression as compared to WT LACV_1978_ (**Figure 6**). Interestingly, across all *Ifn* RNA expression, the S_1978_**M_1960_**L_1978_ reassortant was not significantly different than PBS-infected mice suggesting that the M_1978_ segment plays a major role in inflammation during infection. Between the WT LACV_1960_ and the S_1960_**M_1978_**L_1960_ viruses, the difference in *Ifn-α, Ifn-β,* and *Ifn-γ* RNA expression was less evident, signifying that although the M-segment plays a significant role, the S and L segments may also mediate neuroinflammation (**Figure 7**). Overall, these results suggest that the M-segment significantly mediates type I and type II IFN production in the brain.

**Figure 6:**
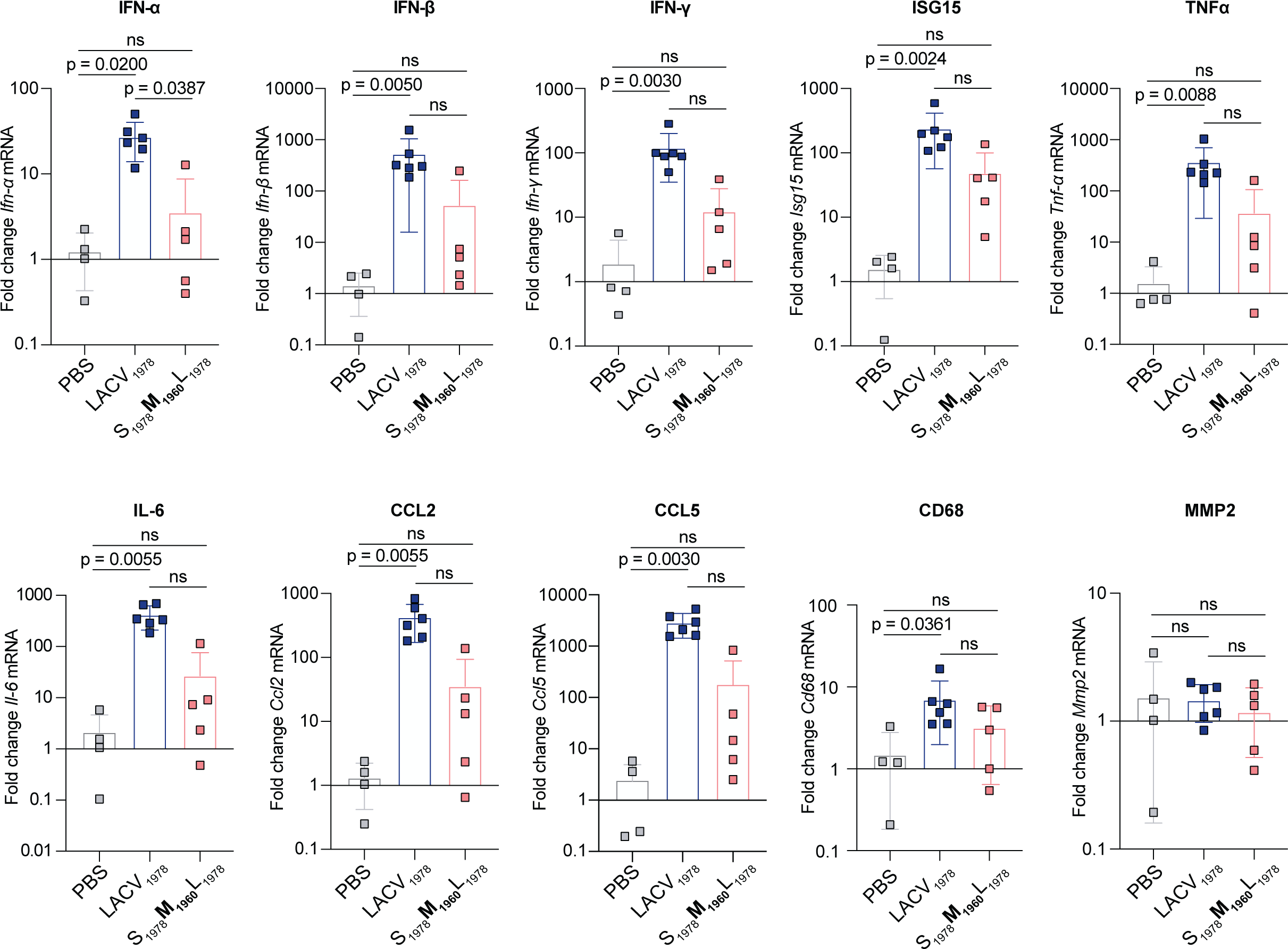
LACV_1960_ M segment decreases IFN and ISG gene expression in the brain. 6-day old C57BL/6J mice were infected with 50 PFU of either LACV_1978_, S_1978_**M_1960_**L_1978_, or mock-infected with PBS. After 3 days post-infection, brains were collected, RNA was extracted and gene expression was quantified via qPCR. Data represent at least two independent trials with n ≥ 5 mice. PBS controls shown are pooled from 5-and 6-day old mice. A Kruskal-Wallis test was performed with Dunn’s multiple comparisons test and p-values are as shown (ns = non-significant). The mean and positive-value standard deviation (SD) are shown for all data.

**Figure 7:**
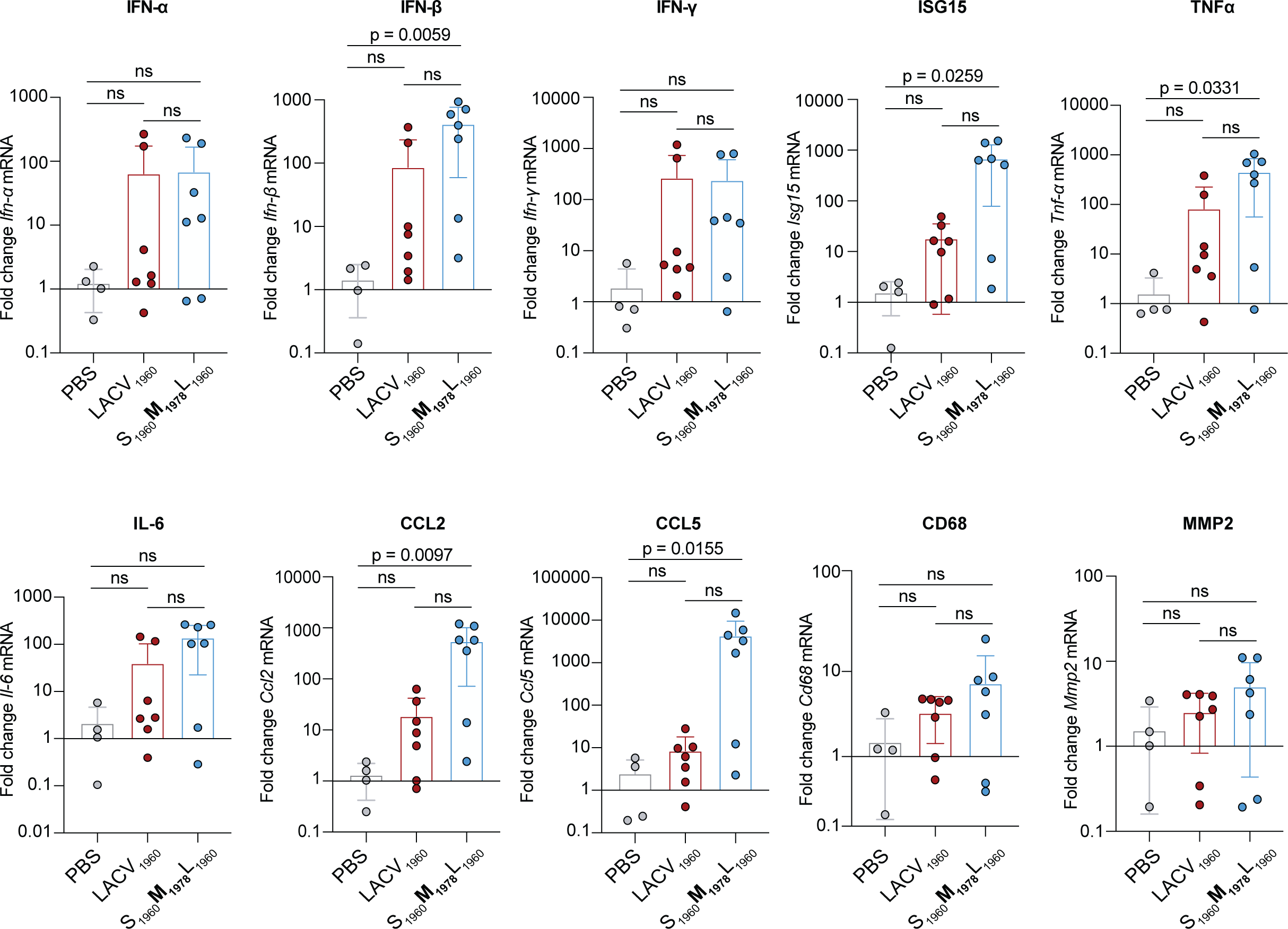
LACV_1978_ M segment enhances IFN and ISG gene expression in the brain. 5-day old C57BL/6J mice were infected with 50 PFU of either LACV_1960_, S_1960_**M_1978_**L_1960_, or mock-infected with PBS. After 3 days post-infection, brains were collected, RNA was extracted and gene expression was quantified via qPCR. Data represent at least two independent trials, with n > 5 mice. PBS controls shown are pooled from 5-and 6-day old mice. A Kruskal-Wallis test was performed with Dunn’s multiple comparisons test and p-values are as shown (ns = non-significant). The mean and positive-value standard deviation (SD) are shown for all data.

We next assessed the effect that IFN production had on the induction of the interferon-stimulated gene (ISG) *Isg15*, pro-inflammatory cytokines (*Tnf-α, Il-6*), and chemokines (*Ccl2, Ccl5*) (**Figure 6 and 7**). Additionally, we evaluated *Cd68* as a marker of microglial activation, which has been shown to be upregulated during inflammation (30), and *Mmp2* expression, a matrix metallopeptidase that regulates blood-brain barrier permeability during infection that has been identified in LACV-infected tissue neuron/astrocyte co-cultures at 48 hours (29). Between WT LACV_1978_ and the S_1978_**M_1960_**L_1978_ reassortant, we observed notable differences for all inflammatory markers and CD68, although not for MMP2 (**Figure 6**). For all markers but MMP2, induction was significantly higher in WT LACV_1978_ infection as compared to PBS, while S_1978_**M_1960_**L_1978_ did not reach a statistically significant difference from PBS. These differences correlate with trends seen in clinical scores, LACV RNA, IFN production, and infectious particles (**Figure 5A-C, and Figure 6**).

Finally, between WT LACV_1960_ and S_1960_**M_1978_**L_1960_, there was an upward trend in induction of *Ifn* RNA expression observable across all IFNs, however only *Ifn-β* had a statistically significant increase (**Figure 7**). We observed an increase in interferon induced inflammatory marker production for S_1960_**M_1978_**L_1960_ for all genes except *Il-6*, *Ccl2*, and *Cd68* (**Figure 7**). Overall, these results illustrate that the M segment reassortants alter both IFN production and inflammatory genes, chemokines, and cytokines, as well as activating microglial cells. Therefore, the M segment may influence neuroinflammation and account for differences in clinical outcomes between reassortants.

## Discussion

Bunyaviruses are significant human pathogens, yet we understand little of the viral determinants of disease. In this project, we aimed to understand how the different orthobunyavirus genomic segments may contribute to replication *in vitro* and *in vivo*. We utilized the orthobunyavirus La Crosse virus (LACV), and generated six reassortants by swapping one segment, either S, M, or L, between two genetically-similar Lineage I WT LACV viruses. These reassortments resulted in one non-rescuable virus and five rescuable viruses that exhibited unique cell-specific phenotypes. Importantly, there are very few amino acid or nucleotide changes between these isolates (**Figure 8**), allowing future work to pinpoint which genomic determinants are driving these phenotypes.

**Figure 8:**
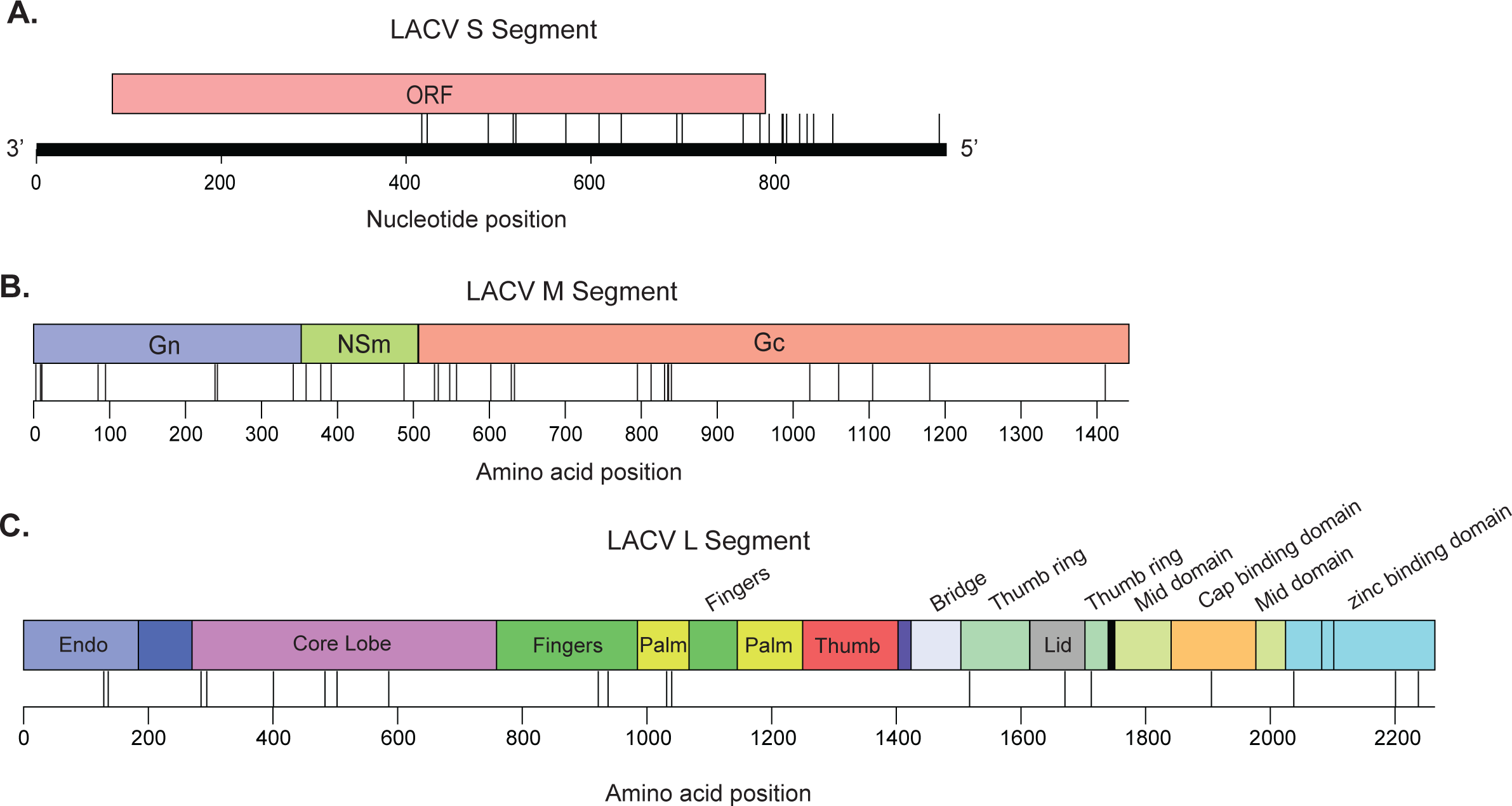
Nucleotide and amino acid differences between LACV strains. (**A**). Schematic of the RNA changes found between the S segments of LACV_1960_ and LACV_1978_. (**B** and **C**) Schematic of the amino acid differences between the M and L protein coding regions of LACV_1960_ and LACV_1978_. Black lines indicate nucleotide (**A**) or amino acid (**B** and **C**) changes. Distinct proteins and domains are indicated.

Swapping the S segment resulted in one non-rescuable virus, **S_1960_**M_1978_L_1978_, while the other reassortant, **S_1978_**M_1960_L_1960_, exhibited a significant growth advantage in both mammalian and insect cell lines. The phenomenon of not being able to rescue one of the reassortants is interesting when considering that both S segments had identical protein sequences indicating a role for the S segment RNA’s interactions with other RNA and/or proteins to be necessary for appropriate replication, packaging and/or egress of the viral particle from the cell (17) (31) (32) (33) (34). Tercero et al. recently elucidated a viral packaging role for antigenomic S RNA in Rift Valley fever virus, a member of the bunyavirus family, which leads us to hypothesize a similar mechanism may be at play in the LACV S segment (35). When we map the nucleotide differences between the two S segments, we find the majority of changes cluster to the 5’ NCR which we hypothesis may play critical roles for packaging and replication (32) (34) (**Figure 8**). While we cannot compare S segment reassortants as is the case of the other segments, the growth advantage of **S_1978_**M_1960_L_1960_ over LACV_1960_ indicates a role for the S segment in increased replication of LACV. Specifically, the S segment encodes a nonstructural protein NSs that has been implicated in suppression of the Type I IFN System which could be aiding in increased virulence in interferon competent NIH/3T3 cells (12).

The M segment reassortants, S_1960_**M_1978_**L_1960_ and S_1978_**M_1960_**L_1978_, had altered growth dynamics in mammalian cells but did not influence virus growth in insect cells when compared to their respective WT parent backbones. One explanation for this phenotype may be that we are using Aag2 *Aedes (Ae.) aegypti* cells and we may see more pronounced phenotypes if *Ae. triseratus* or *Ae. albopictus* cell lines were used. Most interestingly, each M segment reassortant retained the plaque phenotype of their parent WT strain with S_1960_**M_1978_**L_1960_ exhibiting large viral plaques like LACV_1978_ while S_1978_**M_1960_**L_1978_ resulted into small viral plaques like LACV_1960_. The LACV M segment encodes the glycoproteins, Gc and Gn, which are required for viral fusion and entry into the host cell (14–16)(22)(36). Therefore, the LACV M segment likely contains the genomic determinants of viral plaque phenotype formation and dissemination given that the plaque size phenotype remained the same despite changes in S and L segments.

To further investigate if differences in viral plaque and growth phenotypes *in vitro* correlated with viral growth kinetics and neuropathogenesis *in vivo*, we used 5-and 6-day old neonate mice as an *in vivo* model of neuroinvasive LACV infection. The M segment has previously been implicated in eliciting differences in LACV neuroinvasion and pathogenesis (24–26) leading us to question how even minute genomic differences in the M segment could alter neuroinvasion and pathogenesis. When comparing M segment reassortants with their WT backbones, we noted differences in clinical outcomes, viral RNA levels, and viral titers in the brain. When swapping the M_1978_ segment from the more pathogenic LACV_1978_ into the less pathogenic LACV_1960_ backbone, there was an overall increase in disease severity and amount of viral RNA and infectious particles found in the brain. The opposite was true in the S_1978_**M_1960_**L_1978_ reassortant where an attenuative effect occurred and the reassortant was less pathogenic and had lower viral burdens in the brain than the corresponding LACV_1978_ WT. These data support that the M segment is implicated in neurovirulence. Driven by the differences in neuroinvasion, we aimed to identify if the M segment was playing a role in interferon-mediated inflammation. Overall, we noted increases in cellular markers of direct and indirect type I and II cytokine and chemokine induction correlating with the M_1978_ segment, and a general attenuation or lack of induction correlating with the M_1960_ segment. It is interesting that despite no major change in LACV RNA or infectious particles in the brain, significant changes in neuroinflammation are occurring and point to the M segment in influencing neuroinflammation. When we map the amino acid changes between isolates onto the M segment we found changes in all three proteins, Gn, NSm, and Gc (**Figure 8**). Future studies investigating the role of each M segment protein, as well as the remaining S and L proteins, in viral replication and virulence will be essential to understanding what about these segments is driving neuroinflammation.

Finally, the L reassortant, S_1960_M_1960_**L_1978_**, yielded a significant growth advantage over its WT pair, LACV_1960,_ in Vero but not NIH/3T3 or Aag2 cells. There was no significant difference in growth between the inverse pair, S_1978_M_1978_**L_1960_** and LACV_1978_, in either mammalian or insect cells, however, in NIH/3T3, S_1978_M_1978_**L_1960_**showed reduced growth when compared to the WT LACV_1978_. This is notable because it is one of two instances of the pair M_1978_L_1960_ resulting in decreased growth, suggesting this M and L segment combination may be a determinant in levels of viral replication. The LACV_1978_ L segment has a positive replication advantage, while LACV_1960_ L segment has a negligent or attenuative effect on replication when compared to wild-type. Although the two L segments are relatively similar in genetic makeup (**Table 1 and 2**), an unknown difference between these two strains seems to be driving dissimilarities in replication. When we map the amino acid differences between strains onto the L segment, we find changes located in multiple domains critical for polymerase function (18, 23) (**Figure 8C**). Future studies will be dissecting how these changes influence LACV replication *in vitro* and *in vivo*.

Overall, these *in vitro* and *in vivo* studies with all rescuable reassortant viruses concluded LACV replication and virulence are regulated at both the RNA and/or protein level through all three genomic segments. Future research will elaborate on these findings by focusing on what specific molecular features are required for determining these different observed phenotypes in viral growth, plaque size, and virulence, using both *in vivo* models and more pathogenically-relevant cell lines such as a neuron/astrocyte co-culture system that has been shown to be infectible with LACV (29). In case of the S segment, we aim to determine which of the RNA sequence differences between WT strains is contributing to enhancement of viral growth in one instance but complete lack of rescue in the other. We hope that by further interrogating these minute differences in RNA sequence and structure, we will be able to further characterize S segment function in replication and packaging. Taken together, this study exhibits how small molecular changes can determine significant phenotypes in orthobunyaviruses highlighting the need to further characterize the function of each genomic segment in molecular detail.

## Acknowledgements

We thank all members of the Stapleford Lab for helpful discussion on this project. Co-first author order was determined alphabetically. This work was supported by funding from the NYUGSoM Startup and NIAID/NIH R01 AI162774-01A1.

## Materials and Methods

### Cell lines

BHK BSR-T7/5 cells (a gift from Dr. Steven Whitehead at the National Institutes of Health (NIH)), NIH/3T3 cells (gift from Dr. Ken Cadwell at the University of Pennsylvania), and Aag2 *Aedes aegypti* cells (gift from Dr. Paul Turner at Yale University) were grown in complete media comprised of Dulbecco’s Modified Eagle Media (DMEM), 10% fetal bovine serum (FBS: Atlanta Biologicals), 1% nonessential amino acids (NEAA), and 10 mM HEPES (Gibco). BHK BSR-T7/5 cells were maintained under selection with 1 mg/mL Geneticin (Gibco) supplemented in the growth media every other passage. Vero cells (ATCC CCL-81) were maintained in DMEM supplemented with 10% Newborn Calf Serum (NBCS: Sigma). All mammalian cells were maintained at 37°C with 5% CO2, while insect cells were maintained at 28°C with 5% CO2. All cell lines were confirmed mycoplasma free using the Lookout Mycoplasma PCR detection kit (Sigma-Aldrich).

### Viruses

The La Crosse virus (LACV) LACV_1978_ infectious clone system was a gift from Dr. Steven Whitehead at the NIH (21). The LACV_1960_ infectious clone system was a gift from Dr. Friedemann Weber at Justus-Liebig University, Germany (11). Infectious virus was generated by transfecting BHK BSR-T7/5 cells with 2 μg of the S, M, and L segments using the TransIT-LT1transfection reagent (Mirus Bio) following the manufacturer’s protocol. The day after transfection, media was replaced and cells were incubated at 37°C for five days. After incubation, the supernatant was collected and used to inoculate a flask of Vero cells to generate a working stock. Working stocks were harvested when Vero cell cytopathic effect was roughly 70%. Stocks were then centrifuged at 1,200 x g for 5 minutes to clear debris, aliquoted, and stored at −80°C. Viral titers were quantified by plaque assay on Vero cells as described below. Two independent viral stocks were used for all experiments. All virus stocks were confirmed of reassortment via Sanger sequencing.

### Plaque assay

Infectious virus production was quantified by plaque assay. Briefly, virus samples were serially diluted 10-fold in DMEM. 200 μL of dilutions were then added to a monolayer of Vero cells in a 12-well plate and incubated for 1 hour at 37°C. After incubation, DMEM supplemented with 2% FBS and 1% antibiotic-antimycotic was mixed with 0.8% agarose and then overlayed onto the cells for an additional incubation at 37°C. After 4 days, cells were fixed with 4% formalin for 1 hour, after which the agarose plugs were removed and the cells were stained with 0.1% crystal violet in 20% ethanol to visualize plaques. Plaques were counted manually and virus titer was calculated using the lowest countable dilution.

### Growth kinetic assay

Virus growth kinetics were determined via growth assay with multiple time points. Vero, NIH/3T3, or Aag2 cells were plated in 24-well plates about 24 hours prior to infection and incubated at either 37°C (mammalian cells) or 28°C (insect cells). Initial cell densities were as follows: 55,000 Vero cells/well, 50,000 NIH/3T3 cells/well, 200,000 Aag2 cells/well. 100 μL of virus was added to Vero and NIH/3T3 cells at a MOI of 0.1. Aag2 cells received 100 μL of virus at a MOI of 0.03. Infected cells were incubated for 1 hour at 37°C (mammalian) or 28°C (insect). Virus was then removed and cells were washed twice with PBS. 500 μL of complete media was added and cells were then incubated at 37°C (mammalian) or 28°C (insect) for 48 hours. 100 μL of supernatant was collected and replaced with 100 μL fresh complete media at time points 0, 8, 24, 32, and 48 hours post-infection. Supernatant samples were frozen at −80°C. Viral titers were quantified via plaque assay on Vero cells as described above.

### Mouse infections

Mice were maintained in a pathogen-free facility at the NYU Grossman School of Medicine. Individual C57BL/6J (Jackson Laboratory) breeding pairs were established and each litter produced was used as a biological replicate. 5-or 6-day old mice were infected subcutaneously in the back with 50 PFU of each LACV virus; wild-type virus and a paired reassortant virus were matched within each experiment. As an uninfected control, one mouse in each cage was injected with an equal volume of PBS. Mice were then monitored daily and euthanized 3 days post-infection. At time of harvest, a clinical score was noted as indicated in **Table 3** and the brain was collected in DMEM with 2% FBS. The tissue was homogenized for 5 minutes at the setting 12 using a Bullet Blender Storm Pro (Next Advance) and clarified by centrifugation. Viral titers were quantified by plaque assay as described above. All *in vivo* experiments were completed in accordance with the NYU Grossman School of Medicine Institutional Animal Care and Use Committee (IACUC) guidelines (protocol no. IA16-01783).

### RNA extractions and qPCR

For RNA extractions, 400 μL of TRIzol^TM^ reagent (Invitrogen) was added to 100 μL of clarified brain tissue homogenate. RNA extractions were performed following the manufacturer’s guidelines, and extracted RNA was diluted in 400 μL of nuclease-free water. Host gene expression was measured by qPCR using PowerSYBR® Green PCR Master Mix (Thermo Fisher Scientific) with primers in **Table 4** and following manufacturer guidelines. From extracted RNA, cDNA was synthesized with the Maxima H minus-strand kit (Thermo Scientific) using random hexamer primers with the following protocol: 25°C for 10 min, 50°C for 30 min, and 85°C for 5 min. Using a QuantStudio 3 qPCR instrument (Applied Biosystems), cDNA was used in qPCR reactions with the following program: 40 cycles of 95°C for 15 s, 60°C for 1 min. Melt curves were individually analyzed and data collected using QuantStudio software v1.4 (Thermo Fisher Scientific). To determine relative gene expression, each gene was compared to *18S* expression to calculate the ΔCT. To express fold change normalized to PBS-infected controls, the ΔΔCT of each sample was calculated against the average ΔCT of PBS-infected mice. For PBS-infected mice, the ΔCT from 5-and 6-day old mice were pooled.

**Table 4:**
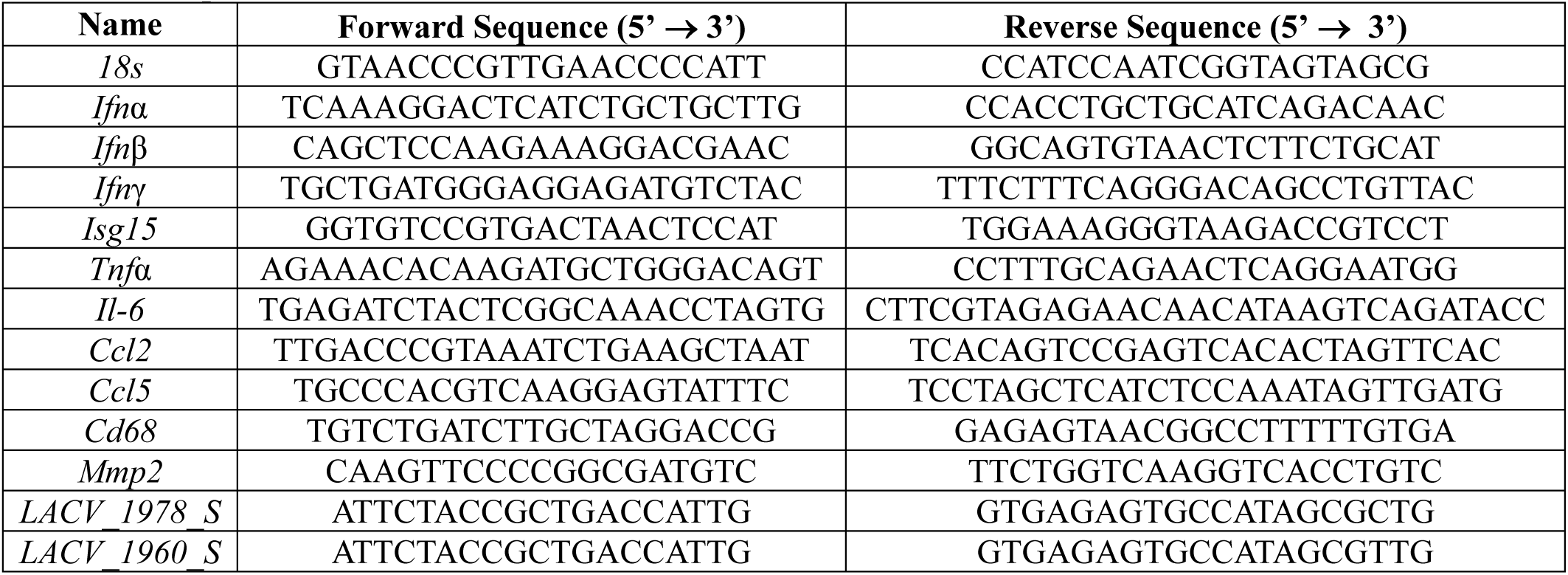
qPCR Primers.

### Protein and nucleotide alignments

RNA conservation within WT strains was aligned using Smitt-Watermen RNA alignment and protein conservation within WT strains was generated using ClustalW Multiple Sequence alignment with MegAlign Pro (DNASTAR; Version 17.4.1.17).

### Sanger sequencing

Viral RNA was extracted from each virus stock and cDNA was synthesized as described above. Each segment of the LACV genome was amplified utilizing primers specified in **Table 5** and a Phusion High-Fidelity PCR Master Mix Kit with HF Buffer (Thermo Scientific). All LACV segments were confirmed by a 1% agarose gel and sent to Sanger sequencing using the primers in **Table 5** at Genewiz (South Plainfield, NJ). LACV samples sent for Sanger sequencing were aligned using SeqMan Ultra (DNASTAR) for verification of reassortment.

**Table 5:**
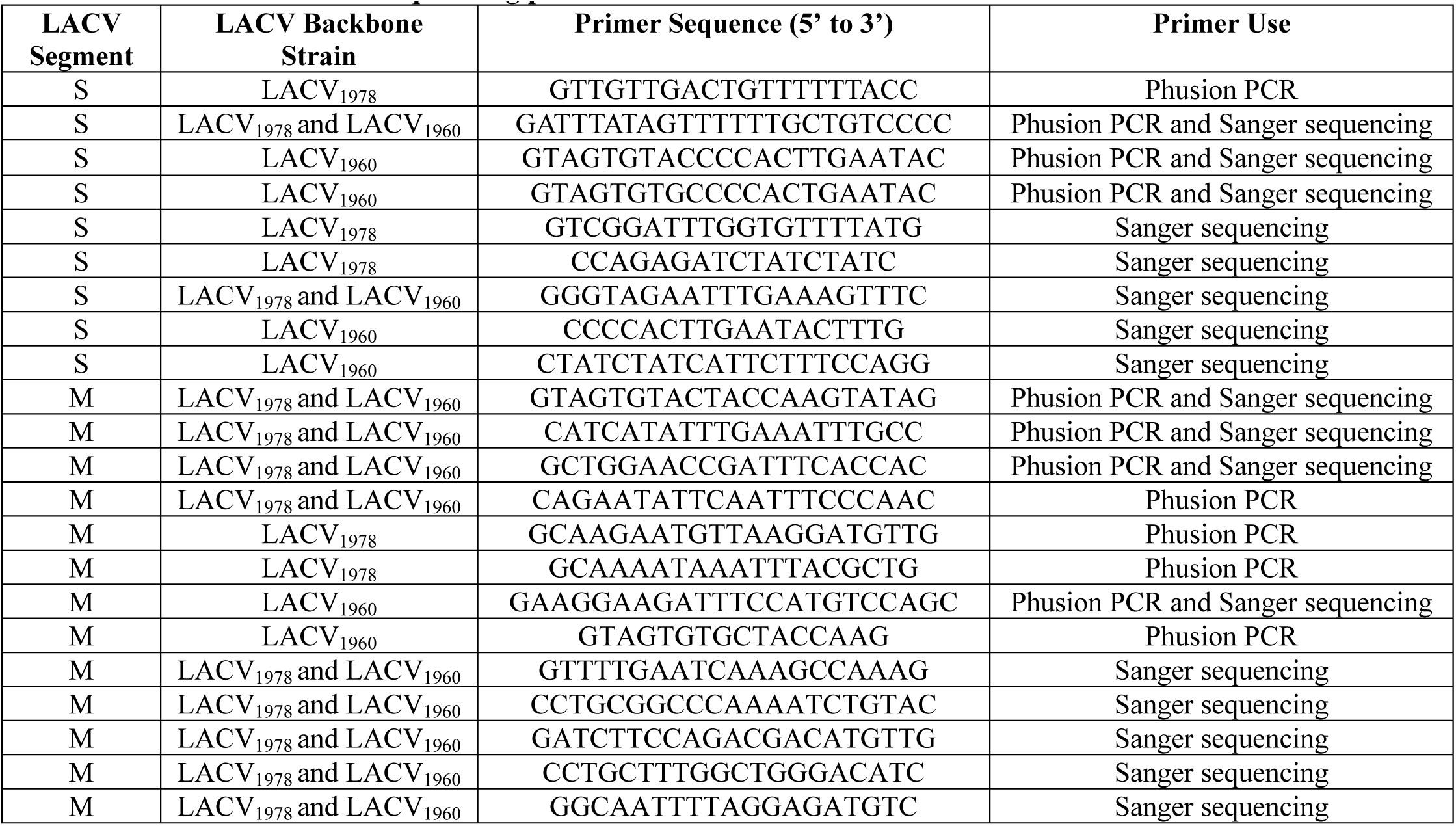
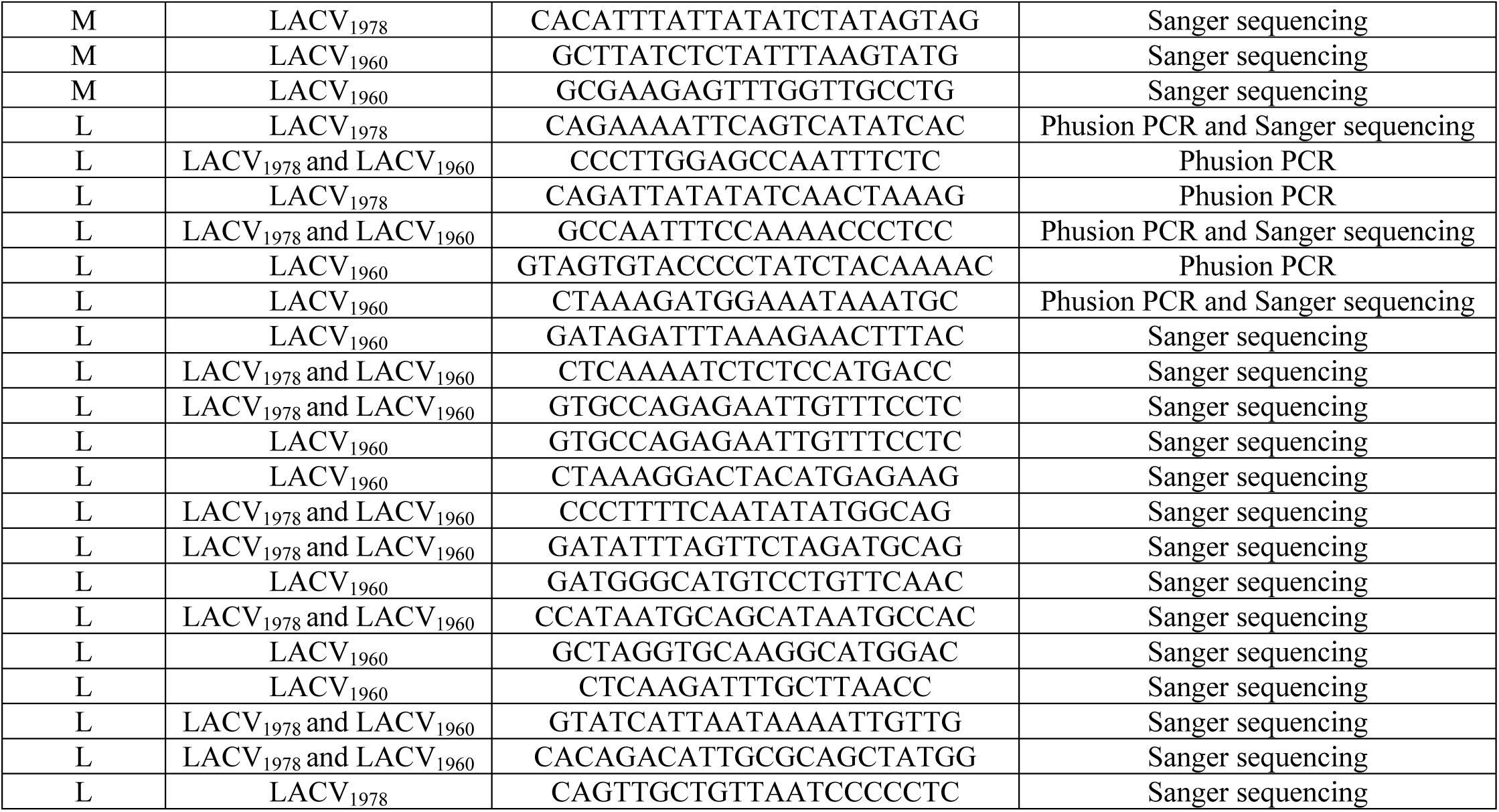
LACV PCR and sequencing primers.

### Statistics and data analysis

All data was analyzed using GraphPad Prism (Version 10). A p-value < 0.05 is considered statistically significant and specific tests are indicated in the Figure Legends. All *in vitro* experiments were completed with at least three biological replicates and internal technical duplicates using two independent stocks of virus. All *in vivo* experiments were completed at least twice and used n ≥ 4 mice for each condition.

